# Low spontaneous mutation rate in a complex multicellular eukaryote with a haploid-diploid life cycle

**DOI:** 10.1101/2022.05.13.491831

**Authors:** Marc Krasovec, Agnieszka P. Lipinska, Susana M. Coelho

## Abstract

The spontaneous mutation rate *µ* is a crucial parameter driving evolution and biodiversity. Mutation rates are highly variable across species, suggesting that *µ* is susceptible to selection and drift and that species life cycle and life history may impact its evolution. In particular, asexual reproduction and haploid selection are expected to affect mutation rate, but very little empirical data is available to test this expectation. Here, we sequence 30 genomes of a parent-offspring pedigree in the brown algae *Ectocarpus* to test the impact of its life cycle on mutation rate. *Ectocarpus* alternates between a haploid and a diploid stage, both multicellular and free living, and utilizes both sexual and asexual reproduction. *Ectocarpus* is therefore a unique model to empirically test expectations of the effect of asexual reproduction and haploid selection on mutation rate evolution. We estimate that *Ectocarpus* has a nucleotide mutation rate of *µ*_*bs*_=4.07×10^−10^ per site per generation, a surprisingly low number for a multicellular complex eukaryote. Effective population size (*N*_*e*_) and genome size could not explain this low mutation rate. We propose that the haploid-diploid life cycle, combined with extensive asexual reproduction may be key drivers of mutation rate.

## Introduction

The spontaneous mutation rate (*µ*) dictates the amount of new genetic diversity generated in a species, population, or genome each generation, and reflects the fidelity of genome transmission. The importance of mutations has been recognized for more than a century, and great efforts have been made to understand the mechanisms underlying mutations and mutation rate evolution (Morgan 1910; Muller 1928; Haldane 1937; Demerec 1937). In the 1960s, Mukai statistically estimated the rate of deleterious mutations in *Drosophila* (Mukai 1964; Keightley and Eyre-Walker 1999) using the decrease of fitness each generation during mutation accumulation experiments (MAE). MAEs, combined with genomic sequencing, remain an accurate and widely used method to directly measure spontaneous mutation rates, using the following of MA lines from a clone (or from an inbred couple in the case of sexual species) under minimal selection (Halligan and Keightley 2009; Lynch *et al*. 2016; Katju and Bergthorsson 2019). Selection is reduced during the experiment by imposing regular bottlenecks, generally at one cell or one couple, to decrease the population size of the MA lines. Such dramatic bottlenecks increase the drift (the random chance of allele frequency variation) to a level where selection is negligible. This allows the fixation of all mutations, including the deleterious and thereby gives access to the total spontaneous mutation rate. However, MAEs are sometimes not feasible because they require the following of dozens of lines during hundreds or thousands of generations. For species with long generation times, a pedigree with parent-offspring genome sequencing is more appropriate and this approach has been employed frequently notably in primates (Pfeifer 2017; Koch *et al*. 2019; Wang *et al*. 2020) and plants (Krasovec *et al*. 2018).

To date, spontaneous mutation rates have been estimated in more than seventy species, including several groups of eukaryotes, bacteria, and archaea (Table S1). We observe a variation of four orders of magnitude between ciliates, with mutation rates of 7.6×10^−12^ (Long *et al*. 2018b), and the great apes, including humans, with mutation rates higher than 1.0×10^−8^ mutations per site per generation (Besenbacher *et al*. 2016, Besenbacher 2019). The “drift barrier” hypothesis has been widely accepted to explain such wide variation (Sung *et al*. 2012a; Lynch *et al*. 2016). According to this hypothesis, the mutation rate per site is lower in species with large effective population size (*N*_*e*_) because selection efficiently favours a small *µ*, which reduces the mutation load due to deleterious mutations. However, in species with a small effective population size, *µ* cannot be maintained at a low rate because of stronger drift which can counteract selection. Another possible contributor to mutation rate variation between species is the effect of genome size. Genome size may impact the cost of replication fidelity in species with large genomes, which is consistent with the positive correlation between genome size and mutation rate (Smeds *et al*. 2016; Krasovec *et al*. 2017). However, a potential confounding factor is that species with large genomes may have small effective population sizes (Lynch and Conery 2003), leading species with large genomes to have a high mutation rate following the drift barrier hypothesis. It is worthwhile to note, however, that the link between genome size and effective population size is debated (Charlesworth and Barton 2004; Whitney and Jr 2010; Lynch 2011; Whitney *et al*. 2011).

Current hypotheses are based on highly phylogenetically biased samples, particularly across eukaryotes. Indeed, the majority (>85%) of the estimates is based on only two eukaryotic groups, Opisthokonta and Archaeplastida (Table S1). Exceptions include one estimate for diatoms (Krasovec *et al*. 2019); four in alveolates (Sung *et al*. 2012b; Long *et al*. 2018b), one in haptophytes (Krasovec *et al*. 2020) and one in Amoebozoa (Saxer *et al*. 2012; Kucukyildirim *et al*. 2020). Interestingly, the mutation rates and spectra of these non-model species are strikingly different when compared to classical plant and animal models. Such a discrepancy may relate to organismal life cycle. For example, the low mutation rates in *Paramecium* (Sung *et al*. 2012b; Long *et al*. 2015) may be explained by the unusual life cycle of ciliates, in which a transcriptionally silent germline genome undergoes rounds of cell division between sexual cycles. Selection would favour a low mutation rate to limit the number of deleterious mutations accumulated in the germline genome before sexual reproduction. In the case of *Dictyostelium discoideum* (Amoebozoa), the short indel mutation rate is higher than that for single nucleotide mutations (Kucukyildirim *et al*. 2020), contrary to observations in model eukaryotes (Sung *et al*. 2016). In the haptophyte *Emiliania huxleyi*, the nucleotide mutation rate from GC to AT is lower than that from AT to GC (Krasovec *et al*. 2020), indicating that its mutational process tends to increase genome GC content, which is also the inverse of most studied eukaryotes.

These cases, as well as results from many other non-model organisms, highlight that the diversity of mutation rates in eukaryotes is underestimated. There is an enormous diversity of genome structures and life cycles across eukaryotes differing from classical biological models that may impact mutation rate evolution. Therefore, increasing the breadth of studied species across the tree of life is of critical importance to generate a more complete view of the causes, consequences and evolution of mutation rates.

Here, we test the effect of a haploid-diploid life cycle that combines both successive asexual generations and the presence of a persistent haploid stage on mutation rate evolution. Several theoretical predictions have suggested that such a life cycle may considerably reduce the mutation rate. First, asexual reproduction or reduced recombination are expected to increase the strength of selection for a low mutation rate because mutator alleles stay linked to a single lineage, increasing mutational load due to the accumulation of deleterious mutations over generations (Kimura 1967). This idea is supported by modelling approaches when drift has limited effect (Gervais and Roze 2017). A lower mutation rate would also be advantageous in this context as an extensive haploid phase is expected to increase susceptibility to deleterious mutations and thus increase the haploid purifying selection efficiency against mutator alleles, contrary to animals with a gamete-limited haploid phase.

To test the prediction that haploid selection during the haploid-diploid life cycle in combination with asexual reproduction decreases the mutation rate, we employed the model brown alga *Ectocarpus. Ectocarpus* is a multicellular complex organism representative of an important, but underexplored, eukaryotic group, the brown algae. Brown algae have been evolving independently from other eukaryotic multicellular groups (plants and animals) for more than a billion years (Coelho and Cock 2020). They represent the third most developmentally complex multicellular lineage on the planet. *Ectocarpus* has a haploid-diploid life cycle alternating between two independent, free living complex multicellular stages which are morphologically distinct: the gametophyte (haploid) and the sporophyte (diploid) (Bothwell *et al*. 2010; Coelho and Cock 2020). In the field, the two stages may inhabit different ecological niches and are often present during different seasons of the year (Couceiro *et al*. 2015), with a considerable portion of their life cycle spent in the haploid phase, potentially subject to haploid purifying selection (Immler and Otto 2018). Moreover, *Ectocarpus* can reproduce both sexually and asexually (through spores produced from diploid sporophytes or through parthenogenesis via non-fertilised gametes which regenerate as haploid individuals), and some populations reproduce almost exclusively asexuality (Couceiro *et al*. 2015). Consequently, *Ectocarpus* supplies an opportunity not only to study mutation rate evolution in a broad taxonomic context, but also to investigate the effect of haploid purifying selection and asexual reproduction.

In this study, we generated extensive genomic sequencing data from a pedigree of *Ectocarpus* (Coelho *et al*. 2012b, 2020; Coelho and Cock 2020) to directly estimate the mutation rate and effective population size. We show that the spontaneous mutation rate of this multicellular organism is remarkably small, on the order of mutation rates of bacteria or unicellular eukaryotes, while its effective population size is on the order of that of other multicellular organisms. We propose that the combination of haploid purifying selection and extensive asexual reproduction during the haploid-diploid life cycle may be the driving forces explaining the unusually low mutation rate of this organism.

## Methods

The *Ectocarpus* sp7 inbred lineage was generated by genetic crosses between siblings over eight meiotic generations from a wild type field collected diploid sporophyte (Ec17; Figure S1). We then started a lineage from this cross with the sporophyte individual Ec372SP, producing successive generations of sporophytes and gametophytes. We sequenced the genome of Ec372SP (Generation 0), two parent individuals from a first asexual reproduction of Ec372SP (419f and 420m), and 30 gametophytes originating from the individual 467SP obtained after the production of 5 generations of gametophytes and 6 generations of sporophytes (Figure S1). DNA was extracted using the OmniPrep Plant kit (G-Biosciences) following the manufacturer’s instructions. The DNAseq libraries were prepared following a PCR-free protocol (Collibri PCR-free PS DNA library prep kit, ThermoFisher) and the genomic DNA of each individual was sequenced by Illumina NovaSeq with 150 bp paired end reads.

Raw reads were trimmed with fastp (Chen *et al*. 2018) and trimgalore (https://www.bioinformatics.babraham.ac.uk/projects/trim_galore/) to remove poly(G) tails and overrepresented sequences (Table S2). Cleaned reads were then mapped onto the reference *Ectocarpus* genome (EctsiV2 from ORCAE database: https://bioinformatics.psb.ugent.be/orcae, (Sterck *et al*. 2012)) assembled with bwa mem (Li and Durbin 2010), and bam files were treated with samtools (Li *et al*. 2009). SNP calling to detect nucleotide and short insertion-deletion mutations was done with HaplotypeCaller from GATK (McKenna *et al*. 2010). Structural mutations were detected with lumpy (Layer *et al*. 2014), delly (Rausch *et al*. 2012) and svaba (Wala *et al*. 2018). Sites were considered callable with a coverage >9x in all samples. Then, *de novo* mutation candidates were filtered as follows: (1) the mutation must be found in only one individual; (2) the alternative allele must be 100% of the coverage of the site; (3) none of the other individuals have any alternative reads even at low quality; (4) the site is haploid in all non-mutated individuals; (5) mutation candidates were manually checked in the pileup file of all individuals; and (6) for structural mutations, candidates at a single position in a single individual were removed if similar variants were found near the position in other individuals. Because we considered only mutations appearing in a single individual, shared mutations appearing from Ec372SP to Ec467SP are not counted in the study.

The mutation rate was calculated as follows: *n*/(*G\**x30x*g*) with *n* = the number of mutations, *G\** = the number of callable sites, 30 = the number of sequenced individuals after 467SP, and *g* = the generation number (*g*=1). Coverage analysis was done with bedtools (Quinlan and Hall 2010). Structural variants were not explored in the scaffold Chr_00, which represents a collection of all contigs that could not be placed on the chromosomes.

To test the putative dosage compensation of duplicated chromosomes in the individual L467_27, we performed transcriptome analysis using triplicate samples of L467_27 with L467_26 as a control. Total RNA was sequenced by Illumina HiSeq with 100 bp paired-end reads. Raw reads were mapped against the reference transcriptome with RSEM (Li and Dewey 2011) with Bowtie2 (Langmead and Salzberg 2012) to get TPM values for each genes. Only genes with TPM higher than 1 were selected for analysis.

## Results

Spontaneous mutation rate We analysed genomic data from the individual at the origin of the lineage (the sporophyte Ec372SP), two parent individuals from the first generation and 30 progenies. Mutations were called on 163,675,306 callable sites corresponding to 83.46% of the genome (Table S3). No structural or short indel mutations were found, but we detected two nucleotide mutations, one from A to G on chromosome 16 at position 1,499,719 (intergenic, individual L467_25) and one from C to A on chromosome 11 at position 645,251 (intron of the gene Ec-11_000650, individual L467_27). This gives a nucleotide mutation rate *µ*_*bs*_=4.07×10^−10^ mutations per site per generation (Table 1), making *Ectocarpus* one of the rare multicellular species with a nucleotide mutation rate below 1.0×10^−09^ (Table S1, Figure 1). The GC content of *Ectocarpus* is 53.59%. Based on the two identified nucleotide mutations, *µ*_*bs GC->AT*_=4.38×10^−10^ and *µ*_*bs AT->GC*_=3.81×10^−10^, giving a modest mutation spectrum bias toward GC to AT mutations of 1.15. However, this result is a preliminary estimate, as it is based on only two mutations. Although we identified no indels or structural mutations, if we assume the occurrence of one hypothetical indel and one hypothetical structural mutation in our pedigree, the total mutation rate would be below 8.15×10^−10^ mutations per site per generation (Table 1).

**Table 1.** Spontaneous mutation rates of Ectocarpus. µ_bs_, nucleotide mutation rate per site; µ_id_, short insertion-deletion mutation rate per site; µ_st_, structural mutation rate per site; U_bs_, nucleotide mutation rate per genome; U_id_, short insertion-deletion mutation rate per genome; U_st_, structural mutation rate per genome. µ_id_, µ_st_, U_id_ and U_st_, are calculated assuming one hypothetical mutation.

| $\mu_{bs}$ | $\mu_{id}$ | $\mu_{st}$ | $\mu_{tot}$ | $U_{bs}$ | $U_{id}$ | $U_{st}$ | $U_{tot}$ |
| --- | --- | --- | --- | --- | --- | --- | --- |
| $4.07 \times 10^{-10}$ | $< 2.04 \times 10^{-10}$ | $< 2.04 \times 10^{-10}$ | $< 8.15 \times 10^{-10}$ | 0.08 | $< 0.0400$ | $< 0.0400$ | $< 0.16$ |

**Figure 1.**
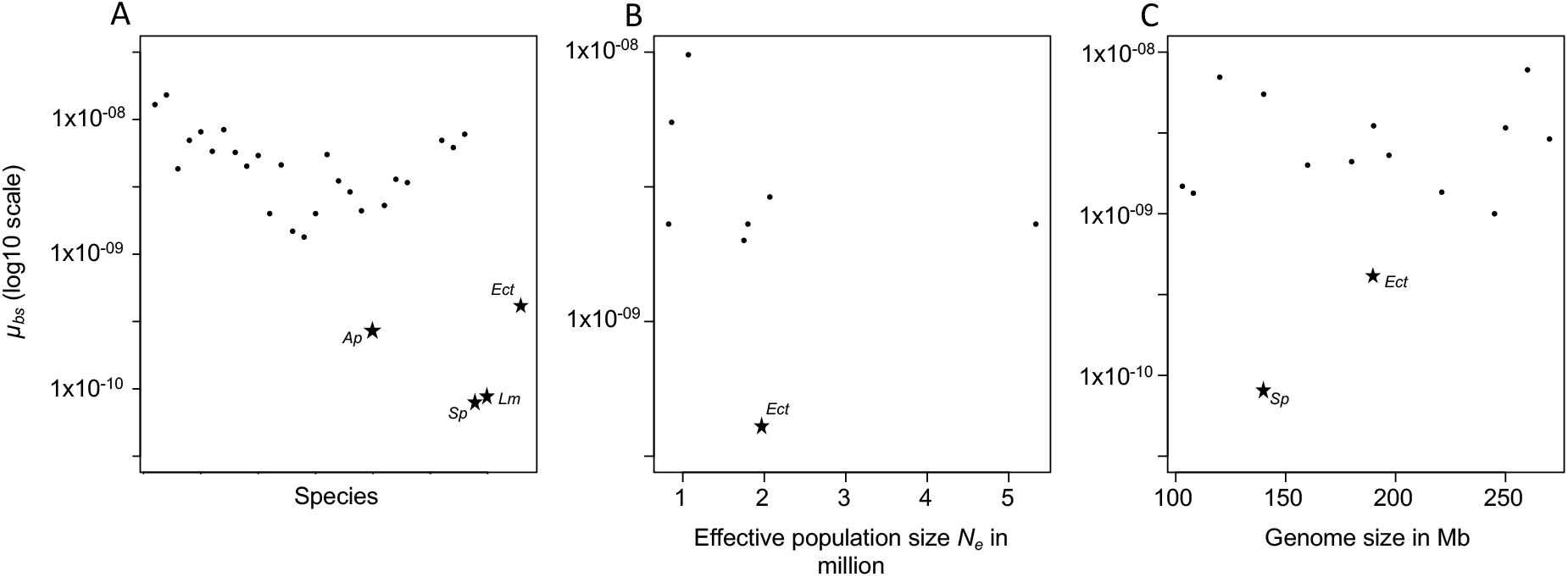
A. Nucleotide mutation rates µ_bs_ of multicellular species (data in Table S1). B. Nucleotide mutation rates µ_bs_ of species with effective population size N_e_ between ∼1 to ∼ 5 million, on the same order as Ectocarpus (N_e_∼2 million). Data from Table S1 of Lynch et al. 2016. C. Nucleotide mutation rates µ_bs_ of species with genome size between ∼100 to ∼ 300 Mb, on the same order as Ectocarpus (214 Mb). Ap: Acyrthosiphon pisum; Sp: Spirodela polyrhiza; Lm: Lemna minor; Ect: Ectocarpus.

### High chromosome duplication rate in *Ectocarpus*

Assessment of raw genome coverage revealed a duplication of 4 chromosomes (C14, C16, C18, C19) in the individual L467_27 (Figure 2). Assuming four independent whole genome duplication (WCD) events, this finding suggests a chromosome duplication rate of 0.0048 duplications per chromosome per generation or 0.1333 chromosome duplications per cell per generation. However, assuming only one independent WCD event (because these duplications were identified in a single progeny), we calculated these rates to instead be 0.0012 duplications per chromosome per generation and 0.0333 chromosome duplications per cell per generation, respectively.

**Figure 2.**
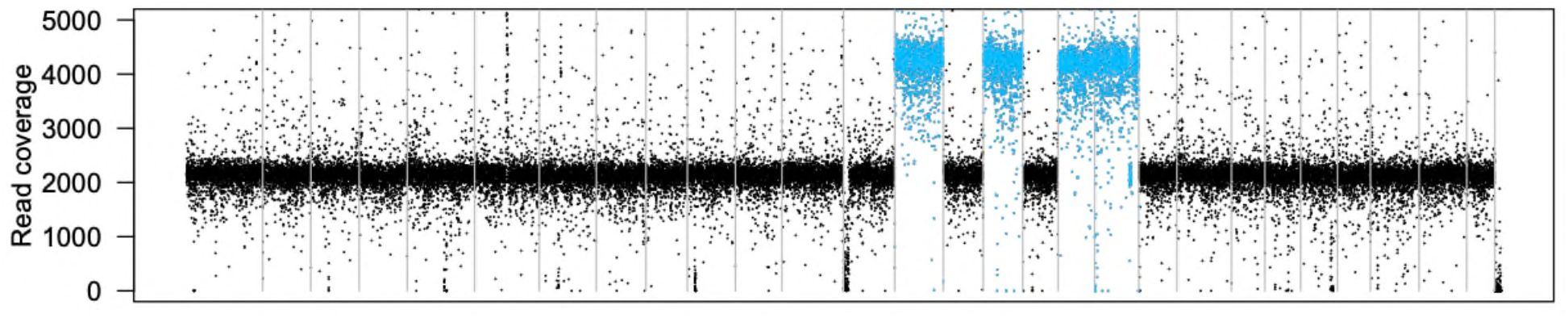
Whole genome raw coverage by 5kb windows showing the four chromosome duplications (in blue) of the L467_27 individual. Grey bars mark chromosome separations.

Chromosome duplications are usually highly deleterious (Sheltzer and Amon 2011), so we investigated if these chromosomal aberrations had an effect on individual fitness. Individual clone L467_27 was cultivated in standard culture conditions (Coelho *et al*. 2012a), and its development was closely followed by regular morphological measures (Table S4) and compared with a sibling without chromosome duplication (L467_26). After 3 weeks in culture, L467_27 exhibited an extensive decrease in fitness, with its growth being markedly slower (Table S4, Figure S2). When cultivated in fertility-inducing conditions, control strain L467_26 produced plurilocular sporangia after 18 days as expected, while L467_27 did not produce meiotic reproductive structures (unilocular sporangia) even after 25 days. Taken together, these results suggest that the identified chromosome duplications did have a negative effect on the growth and reproductive fitness of this clone. To explore the effect of the WCDs on transcription, we measured the level of genome-wide transcription of L467_27 and its sibling control (L467_26). Comparative transcriptomic analyses indicated a ∼1.71-fold higher mRNA level for genes located within the duplicated regions compared to non-duplicated regions (Figure S3), suggesting a general lack of dosage compensation in the line harbouring WCDs.

### Effective population size

Our estimate of the spontaneous mutation rate allows us to calculate the effective population size (*N*_*e*_), following *π*_*s*_=4.*N*_*e*_.*µ*. The neutral diversity (*π*_*s*_) of *Ectocarpus* has been previously estimated based on 49 sporophytes from three European and one South American population and six gametophytes from two European and one South American population (Avia *et al*. 2018). The neutral diversity of autosomal genes is *π*_*s*_=0.00323, which leads to an estimated effective population size of 1,984,029. *Ectocarpus* has a haploid UV sex chromosome system (Ahmed *et al*. 2014; Coelho *et al*. 2018, 2019), making it possible to calculate the effective population size of the different parts of the sex chromosome: π_s_ of the pseudo-autosomal region (PAR, the recombining part of the sex chromosome) was estimated to be 0.00439, giving an effective population size of 2,696,560; and π_s_ of the sex-determining region (SDR, non-recombining) is 0.00221, leading to an effective population size of 1,357,494. The higher *N*_*e*_ of the PAR region may be caused by balancing selection between male and female alleles (Avia *et al*. 2018), whereas the smaller *N*_*e*_ of the SDR is likely caused by the absence of recombination (Ahmed *et al*. 2014). For these reasons, we will consider hereafter the effective population size of *Ectocarpus* as the effective population size calculated for autosomal genes. The time of suppression of recombination on the SDR is given following *π*_*s*_=2.*t*.*µ*, where *t* is the divergence time, calculated here as 2,714,988 generations. This result is consistent with an old UV sexual system, as has been previously suggested (Lipinska *et al*. 2017).

## Discussion

### Low level of spontaneous mutation in *Ectocarpus*

Although *Ectocarpus* is a complex multicellular eukaryotic species, our results using pedigree sequencing revealed an exceptionally low spontaneous mutation rate, on the same order of magnitude as that of unicellular organisms such as bacteria (Long *et al*. 2018a), yeast (Lynch *et al*. 2008; Zhu *et al*. 2014) and phytoplankton (Ness *et al*. 2012; Krasovec *et al*. 2017, 2019, 2020). To date, estimates of mutation rates in multicellular species such as insects (Schrider *et al*. 2013; Keightley *et al*. 2014a; b; Liu *et al*. 2017; Oppold and Pfenninger 2017; Krasovec 2021), small vertebrates (Uchimura *et al*. 2015; Smeds *et al*. 2016; Feng *et al*. 2017; Malinsky *et al*. 2018), and plants (Ossowski *et al*. 2010; Xie *et al*. 2016; Krasovec *et al*. 2018) are at least one order of magnitude higher. *Ectocarpus* is therefore one of the few cases of a multicellular species with a very low spontaneous mutation rate (Figure 2A). This observation suggests that the selection pressure for a low mutation rate in *Ectocarpus* may be stronger than for most multicellular species.

### N_e_ and genome size do not explain low µ_bs_ in *Ectocarpus*

Following the drift barrier hypothesis (Sung *et al*. 2012a; Lynch *et al*. 2016), effective population size is key to understand mutation rate variation between species. The effective population size of *Ectocarpus* is about 2 million, a value unlikely to be sufficient to explain its low mutation rate when compared to the available literature (Lynch *et al*. 2016). Species with an effective population size on the same order of magnitude as *Ectocarpus* typically have a mutation rate an order of magnitude higher (Figure 2B). For example, the effective population sizes of *Daphnia pulex, Drosophila melanogaster, Mesoplasma florum, Pristionchus pacificus, Neurospora crassa, Heliconius melpomene* and *Trypanosoma brucei* are between 1 to 5 million, with spontaneous mutation rates from 1.4×10^−9^ to 9.8×10^−9^ mutations per site per generation. Species with a mutation rate similar to *Ectocarpus* instead have a much larger effective population size of 10 million or more (see Table S1, Lynch *et al*. 2016). In other words, the mutation rate of *Ectocarpus* does not fit classical species expectations.

A second characteristic expected to affect mutation rate is genome size, but similarly as with effective population size, multicellular species with similar genome sizes to *Ectocarpus* typically have spontaneous mutation rates one order higher (Table S1, Figure 2C). For example, *Prunus persica* (Xie *et al*. 2016), *Drosophila pseudoobscura* (Krasovec 2021), *Drosophila melanogaster* (Schrider et al. 2013; Keightley et al. 2014a), *Chironomus riparius* (Oppold and Pfenninger 2017), *Daphnia pulex* (Flynn et al. 2016) or *Apis mellifera* (Liu et al. 2017) all have genome sizes between 100 and 300 Mb but substantially higher mutation rates than *Ectocarpus*. The duckweed species *Spirodela polyrhiza* is uniquely similar to *Ectocarpus* in this regard, with a genome size of 140 Mb and a very low mutation rate (Sandler *et al*. 2020).

### Asexuality and persistent haploid stage may explain small µ_bs_ in *Ectocarpus*

The low mutation rate of *Ectocarpus* is unlikely to be explained by effective population size and genome size. Instead, one possible explanation may be this species’ capacity for asexual reproduction. This hypothesis would be in line with the Kimura prediction (Kimura 1967), suggesting that the selection coefficient *k* for a low mutation rate increases when the recombination rate *r* decreases (*r* = 0 for asexual reproduction), given the general assumption of mutations as deleterious. Note that only two other examples of multicellular species with particularly low mutation rates have so far been described: the pea aphids (*Acyrthosiphon pisum)* (Fazalova and Nevado 2020) and the duck weeds (*Lemna minor*) (Xu *et al*. 2019; Sandler *et al*. 2020). These organisms can also reproduce asexually (by budding and parthenogenesis) for several generations before engaging in sexual reproduction, supporting the hypothesis that asexual reproduction indeed may drive the evolution of low mutation rates. However, it appears that although *Ectocarpus* is capable of asexual reproduction, it may be highly sexual in the field, and significant variation of asexual reproduction frequency between populations has been observed (Couceiro *et al*. 2015). Therefore, it is possible that asexual reproduction alone may not fully provide an explanation for the very low mutation rates we observe in *Ectocarpus*.

An additional, non-mutually exclusive explanation for the low mutation rate we observe is the haploid-diploid life cycle of *Ectocarpus*, which includes a persistent, complex, multicellular haploid stage. The gametophyte haploid stage of *Ectocarpus* has a relatively complex morphology, is free-living and macroscopic, persists for several months, and is associated with the expression of the majority of organismal genes (Lipinska *et al*. 2015). In contrast, the haploid stage of animals such as *Drosophila* is limited to the gametes, and in the more common diploid state, genetic dominance has the potential to mask the effects of mildly deleterious mutations. Thus, in *Ectocarpus*, haploid purifying selection may optimise selection against mutator alleles or any *de novo* mutations with effects on mutation rate (Kimura 1967).

### Whole chromosome duplication rate

Whole chromosome duplications have been reported in several mutation accumulation studies in unicellular green algae (Krasovec *et al*. 2022) and yeast (Zhu *et al*. 2014; Liu and Zhang 2019), where the whole chromosome duplication rate per chromosome per generation was below 0.0001. The chromosome duplication rate we measured for *Ectocarpus* was about 100-fold higher than in these species, in striking contrast to its low nucleotide mutation rate. The mutational processes between WCD and point mutations are very different: point mutations result from replication errors, whereas WCDs result from mis-segregation of chromosomes. WCD may result from a mis-segregation of the chromosome set between daughter cells. Aneuploid karyotypes are broadly considered to be deleterious because they cause an imbalance in gene dosage and transcript production (Hou *et al*. 2018). Indeed, the deleterious effect of dosage imbalance can also apply to single gene duplications. In *Caenorhabditis elegans*, excess transcripts from duplicated genes have been shown to be highly deleterious, and selection acts more strongly against duplication of highly transcribed genes (Konrad *et al*. 2018). Mechanisms of dosage compensation, well known for sex chromosomes, may reduce the deleterious effects of such gene dosage imbalance by adjusting the transcription of genes on aneuploid chromosomes (Disteche 2012). However, the relevance of dosage compensation mechanisms for autosomal chromosomes, particularly immediately after duplication events is poorly understood, especially in emergent models such as *Ectocarpus*. In this study, the reduced fitness of individuals with duplicated chromosomes could reflect highly deleterious effects of WCD and lack of dosage compensation, which was further supported by our transcriptomic analysis.

### Structural mutation rate µ_st_ is poorly understood

Understanding of the rate of structural mutations (sub-chromosomal copy number variation<u>s</u>, inversions, large insertion-deletions and chromosomal rearrangements) is comparatively rare in the literature (Katju and Bergthorsson 2019). Spontaneous structural mutation rates have been calculated for only few species such as green algae (Leger-Pigout and Krasovec 2022), yeast (Lynch *et al*. 2008; Zhu *et al*. 2014; Liu and Zhang 2019), *Daphnia* (Keith *et al*. 2016; Ho and Schaack 2021) and *Caenorhabditis elegans* (Konrad *et al*. 2018). Structural mutations are evolutionarily important as they impact a large portion of the genome and may originate major adaptations. Notably, inversions can cause recombination arrest and lead to the creation not only of supergenes (Taylor and Campagna 2016) but may also contribute to the emergence of the non-recombining regions of sex-chromosomes (Charlesworth 2016; Graves 2016). It is therefore crucial to estimate the structural mutation rate to have a complete view of genome dynamics, with the total mutation rate *µ*_*tot*_=*µ*_*bs*_+*µ*_*id*_+*µ*_*st*_. Almost all current estimates include only spontaneous nucleotide and short indel mutations, preventing us from fully understanding mutation spectrum evolution and the total amount of genetic diversity generated each generation.

Together, our work suggests that organismal life cycle plays a significant role in mutation rate evolution and corroborates the predicted effects of asexuality and a persistent complex haploid phase on mutation rate. More broadly, investigations of non-model or emergent model species will significantly improve our understanding of mutation rate evolution across the tree of life.

## Author contributions

Original idea: MK, SMC; Data analysis: MK, Interpretation: MK, AL, SMC; First draft: MK; Final draft, editing and revisions: MK, SMC; Project management and funding: SMC.

## Acknowledgements

We are grateful to Akira Peters for generating the inbred lines, Andrea Belkacemi and Min Zheng for help with DNA and RNA extractions and library preparations, Dorothee Koch for help with algal cultures, and Denis Roze for comments on the manuscript. This work was founded by the Max Planck Society and a European Research Council grant 864038 to SMC.

## Data availability statement

Genomic raw reads are available under the bioproject xxx.

## Supplemental Tables Legends

Table S1. Published nucleotide mutation rates from pedigree and mutation accumulation studies.

Table S2. Genomic coverage and fastq statistics.

Table S3. Number of callable sites with a minimum coverage threshold of >9× per line.

Table S4. Fitness measurements of wild type and lines showing chromosome duplications

## Supplemental Figures

**Figure S1.**
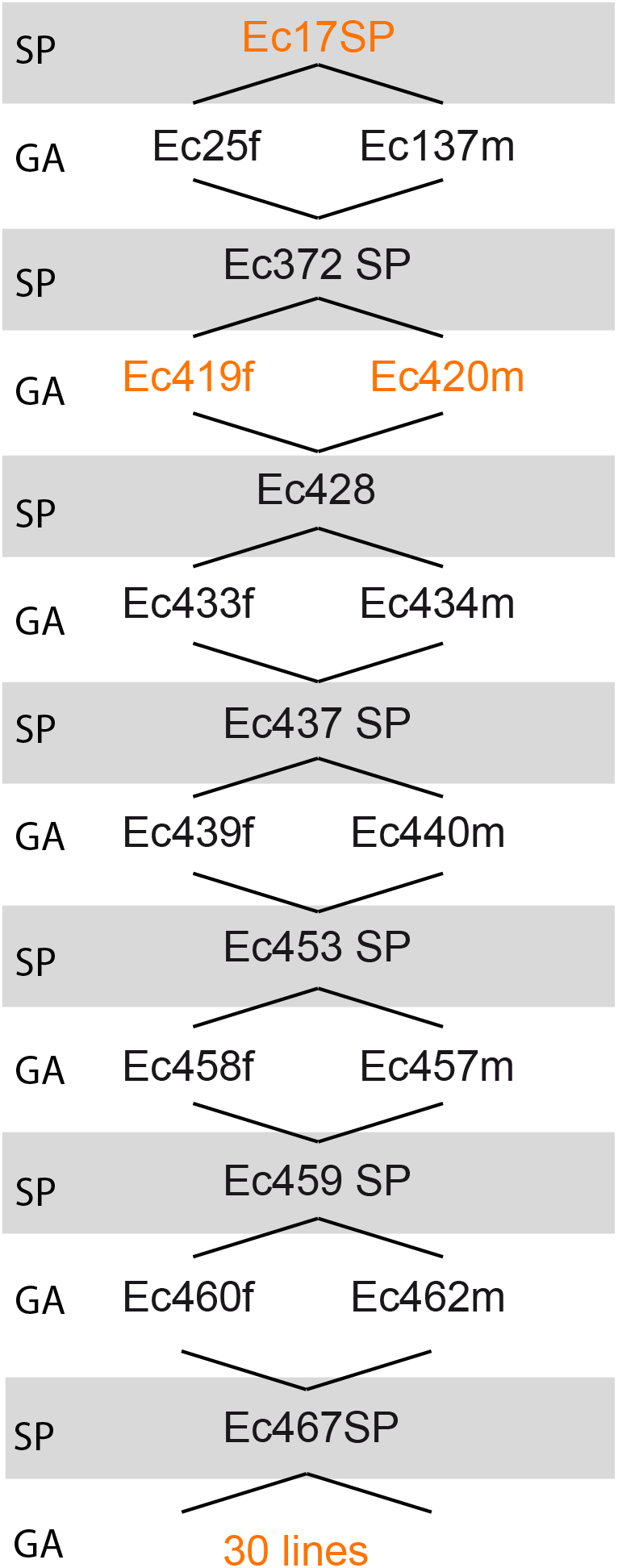
*Ectocarpus* pedigree used in this study. All individuals originate from 372SP. Successive generations led to the individuals 467SP for which we sequenced thirty progenies. Orange: sequenced individuals; GA: gametophyte, haploid; SP: sporophyte, diploid.

**Figure S2.**
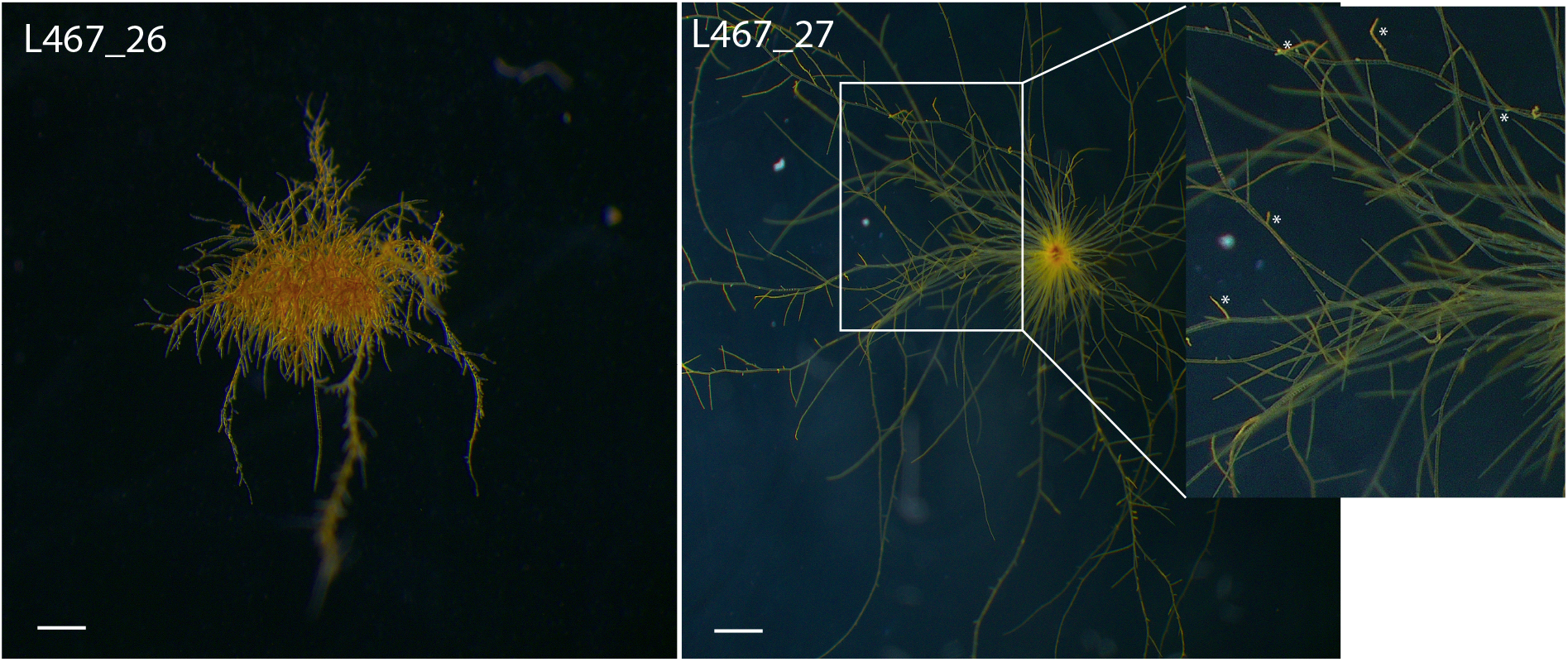
Representative microscopy images of lines L467_26 and L467_27 at day 18 after germination. Asterisks represent plurilocular sporangia (where spores are produced). Sale bar=100µm.

**Figure S3.**
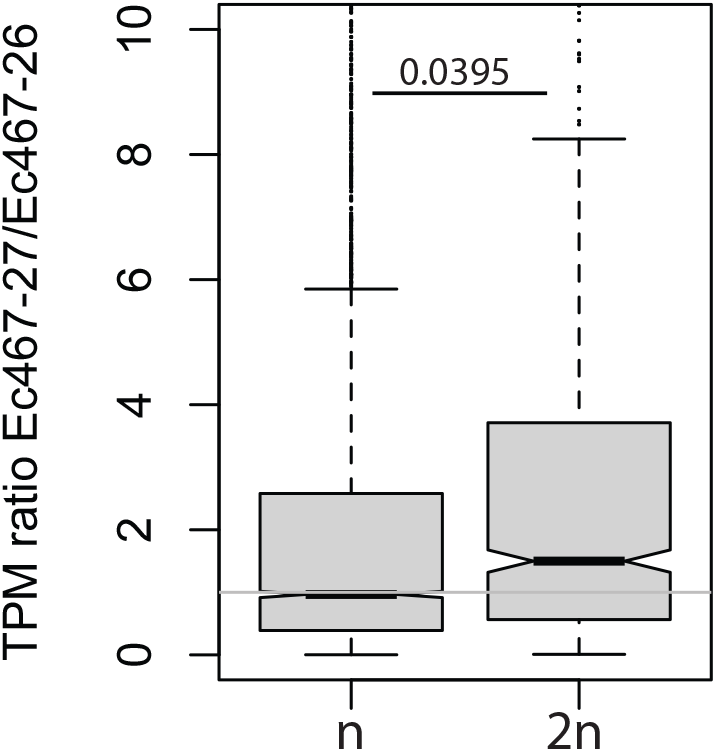
Transcription ratio of genes linked to non-duplicated (n) and duplicated (2n) chromosomes of E467_27. Student test, p-value= 0.0395.

